# Cytokinin overcomes spikelet-driven inhibition of tillering in wheat and barley by delaying meristem development

**DOI:** 10.1101/2025.06.02.657399

**Authors:** Alex Wakeman, Tom Bennett

## Abstract

Correlative inhibition mechanisms allow plants to regulate the number of organs they produce, but act as a developmental limitation on crop yields. Previously, trade-offs have been observed in wheat and barley between the size of individual compound inflorescences (spikes) and the total number of spikes produced by each plant. We aimed to test the hypothesis that these trade-offs represent an uncharacterized form of correlative inhibition in grass reproductive architecture. We show that ‘high spikelet’ lines with spikes containing enhanced numbers of spikelets have reduced tillering and retarded development of later-initiated spikes. Moreover, we show that this effect is driven by the presence of developing, but not mature, spikelets in early-initiated spikes, thus representing a form of correlative inhibition. We show that cytokinin treatment delays the development of spike meristems and formation of spikelets, and as such, can promote the development of additional spikes in high spikelet lines, by providing a longer window for tillering to occur. Our results thus identify cytokinin as a potential target for breeding crops with enhanced yield. Furthermore, our results provide a developmental basis for previously observed yield trade-offs in cereal crops, and illustrate the importance of understanding developmental timing when attempting to breed higher-yielding crops.

## INTRODUCTION

Increasing global population and the effects of climate change on agriculture are culminating in a need to produce increased crop yields whilst simultaneously reducing the quantities of fertiliser, water and arable land we use to grow crops (Cassman *et al*., 2003; Alexandratos and Bruinsma, 2012; Godfray, Poore and Ritchie, 2024). Crop plants are certainly capable of producing far more yield than they do naturally, without increasing resource availability. For instance, crop modelling suggests that in the UK, yields of 21 tons/hectare should be possible with available resources, but typical yields are 8 tonnes/hectare (Mitchell and Sheehy, 2018). There are multiple structural and developmental reasons for this, all of which reflect the fact that plants have evolved to maximise their long-term survival in an uncertain environment, rather than to simply produce as much seed as possible. Plants are therefore inherently ‘cautious’ about resource use, and this is not easily bred out of crop species. Structural limitations include the need to produce additional vegetative tissues to support increased reproductive effort, the need to produce additional reproductive structures to bear seed, the limited time window in which to grow, and the requirement for simultaneous seed-set and maturation for mechanical harvesting. Developmental limitations include the constraints imposed by growth ‘decisions’ made early in development in response to environmental conditions at that time, which will not reflect conditions when seed-set occurs, and the tendency of plants to limit their own growth in response to the growth of existing organs (Walker and Bennett, 2018). This is the phenomenon of ‘correlative inhibition’, the suppression of the formation or growth of new organs by existing structures (Bangerth, 1989).

The most well-known form of correlative inhibition is apical dominance, in which actively growing shoot branches repress the growth of subtending axillary buds; removal of a dominant branch will allow growth of the repressed buds (Domagalska and Leyser, 2011). However, a series of other correlative inhibition relationships exists between different plant structures, with seed acting as particularly strong source of correlative inhibition (Bangerth, 1989; Walker, Wheeldon and Bennett, 2021). Two classes of model, which are not mutually exclusive, have been proposed to explain correlative inhibition. The first is the ‘nutrient diversion’ model, in which organs formed early in development inhibit the formation of later organs by monopolizing the supply of soil-derived mineral nutrients and photosynthetically derived sugars (Phillips, 1975; Cline, 1991). While such models are plausible, they essentially assume that plants only stop initiating organs when nutrients become limiting, which is demonstrably not the case (Walker, Wheeldon and Bennett, 2021). The second class of model, which is rooted in the study of phytohormonal signalling, suggests that existing organs actively release signals that inhibit the development of new organs, thus preventing plants from over-committing resources early in development that will be needed for the development of later structures (Walker and Bennett, 2018). In particular, correlative inhibition has long been associated with the hormones auxin and cytokinin. In the context of shoot branching, auxin is certainly exported from actively growing organs through rootward polar auxin transport, and is certainly able to indirectly inhibit the growth of new branches (Domagalska and Leyser, 2011). Auxin export from seed and fertilized fruit has also been long associated with the ability of older fruit/seed to inhibit the development of new fruit/seed (Bangerth, 1989; Lenser *et al*., 2018; Walker and Bennett, 2018; Haim *et al*., 2021; Sadka *et al*., 2023), or other reproductive structures (Ware *et al*., 2020). On the other hand, cytokinin seems to promote the formation of new organs by overcoming the inhibitory effect of auxin (Müller *et al*., 2015; Lenser *et al*., 2018). Despite the central role of correlative inhibition in the self-limitation of plant growth, and its implications for understanding developmental limits in crop plants, these phenomena remain relatively understudied, particularly in cereals, and poorly characterized at a molecular and developmental level.

Cereal crops of the grass family (*Poaceae*) are a particularly important target in which to realize significant yield increases, since wheat, rice, maize, sugarcane and barley alone constitute more than half of global calorie intake (Ross-Ibarra, Morrell and Gaut, 2007). As members of the same family, cereals share a common shoot and reproductive architecture. During the vegetative period, the primary shoot meristems in grasses initiates a series of leaves and associated axillary meristems, some of which will give rise to vegetative branches (tillers). At the initiation of the reproductive phase, vegetative shoot meristems convert into reproductive inflorescence meristems, some proportion of which will fully develop into a compound inflorescence borne on an elongated stem; in wheat and barley, this is typically referred to as a spike or ear (Koppolu and Schnurbusch, 2019). The primary spike meristem initiates a series of nodes that give rise to secondary inflorescences (spikelets). In barley, each spikelet node develops into a triple spikelet meristem with two lateral and one central spikelet meristem. In most commercial ‘two-rowed’ varieties, only the central spikelet meristem actually gives rise to a spikelet, but in some commercial varieties, all three meristems give rise to spikelets and a ‘six-rowed’ spike (Harlan, 1979; Lundqvist and Lundqvist, 1987; Bellucci *et al*., 2013). In wheat, additional ‘paired’ spikelets can form from a node already bearing a spikelet, like from accessory axillary meristems (Boden *et al*., 2015). Spikelet meristems initiate a series of nodes that develop into floret meristems, which ultimately give rise to florets and seed. Seed yield in grasses therefore reflects the hierarchical and spatio-temporal production of shoots (tillers), primary inflorescences (spikes), secondary inflorescences (spikelets) and flowers (florets) during reproductive development.

There is obvious interest in manipulating the cereal reproductive architecture to increase yields. However, these efforts are confounded by the tendency of yield components to trade off against each other (Sadras, 2007; Reynolds *et al*., 2009; Wolde, Mascher and Schnurbusch, 2019; Abbai *et al*., 2024). Pushing the development of one component typically results in a reduction of other components, with little or no increase in final yield (Abbai *et al*., 2024; Vicentin, Canales and Calderini, 2024). One such example is six-rowed barley, which can result from mutation of any of the *SIX ROWED SPIKE*/*VULGARE ROWED SPIKE* (*VRS*) genes *VRS1*-*VRS5* in barley (Zwirek et al, 2019). *VRS1* encodes an HD-ZIP I transcription factor (Komatsuda et al, 2007), which regulates spikelet fertility downstream of VRS3, VRS4 and VRS5 (Zwirek et al, 2019). VRS5 is a TCP class II transcription factor, and a close homologue of the branching regulatory genes *TEOSINTE BRANCHED1* (*TB1*) in maize and wheat, *FINE CULM1* (*FC1*) in rice, and *BRANCHED1* (*BRC1*) in Arabidopsis (Ramsay et al, 2011). Six-rowed barley is of interest to breeders due to the increased numbers of spikelets per spike, but which is confounded by producing reduced number of tilers and therefore spikes compared to 2-row lines (Zwirek, Waugh and McKim, 2019). In wheat, a *highly branched* (*hb*) mutant which produces supernumerary ‘paired’ spikelets (likely derived from accessory meristems subtending regular spikelets) was identified from a near-isogenic line (NIL) (Dixon *et al*., 2018) derived from the four-parent MAGIC line 0053 (Huang *et al*., 2012). This line was found to exhibit increased *TaTB-D1* transcripts in the inflorescence and tiller buds, resulting in a significantly increased number of spikelets per spike relative to wild type (Dixon *et al*., 2018). However, as with six-rowed barley, this ‘high spikelet’ lines exhibited a significantly reduced tiller number compared to their wild-type background line (Dixon *et al*., 2018). We hypothesised that the *vrs* and *hb* mutants represent examples of correlative inhibition between different components of reproductive architecture in cereals, in which the presence of additional spikelets per spike leads to inhibition of tillering. In this study, using *vrs1* and *hb* mutants as model ‘high spikelet’ lines, we aimed to test this idea, to identify the developmental basis of the phenomena, and to understand how such self-limiting growth might be overcome.

## RESULTS

### Wheat landraces exhibit negative correlation between spikelets and tillers

To assess whether correlative inhibition-type feedbacks might exist within the reproductive architecture of grass species, we examined a panel of 114 spring wheat landraces and elite lines drawn from the CIMMYT, Watkins, Prague and IBTI collections. We assessed a range of shoot architectural traits, including tiller number (at 6, 9 and 12 weeks post germination), final spike number, spikelet nodes per spike, total spikelet nodes, total fertile spikelets, final height, and main tiller/stem diameter. Comparing these traits between lines showed some interesting correlations (or lack thereof) between different components of architecture. In particular, we observed a clear negative correlation between the mean number of spikelets per spike and the number of tillers and spikes produced by each line (Figure 1). This is indicative of a trade-off between the production of new axes of growth (tillers) and the enhanced development of the existing axes (spikelets per spike).

**Figure 1:**
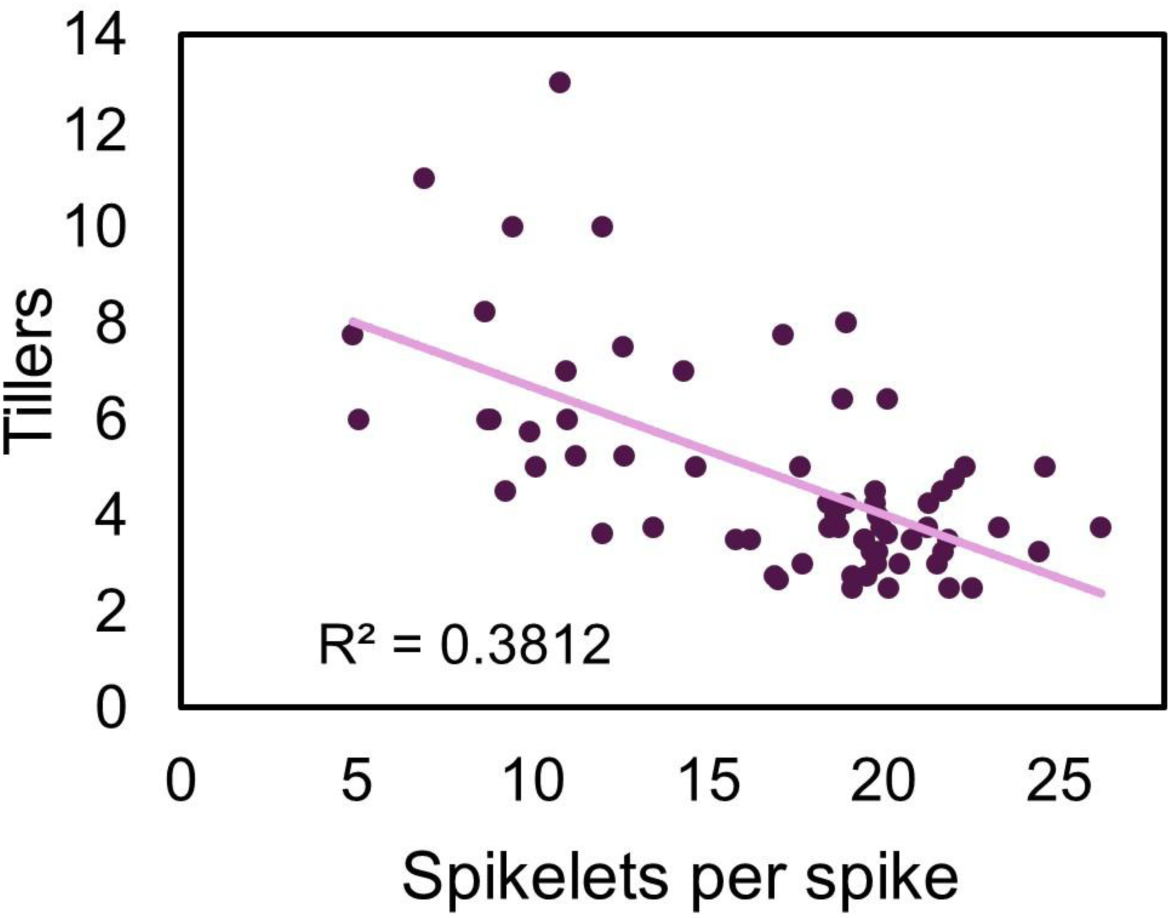
Negative correlation between spikelets per spike and tillers in wheat landraces. Scatterplot showing correlation between mean number of tillers per plant and mean number of spikelets per spike after 12 weeks of growth in 66 spring wheat landraces (n=3-5 per line, R^2^ = 0.38). Each point represents the mean values for an individual landrace

**Supplementary Figure 1:**
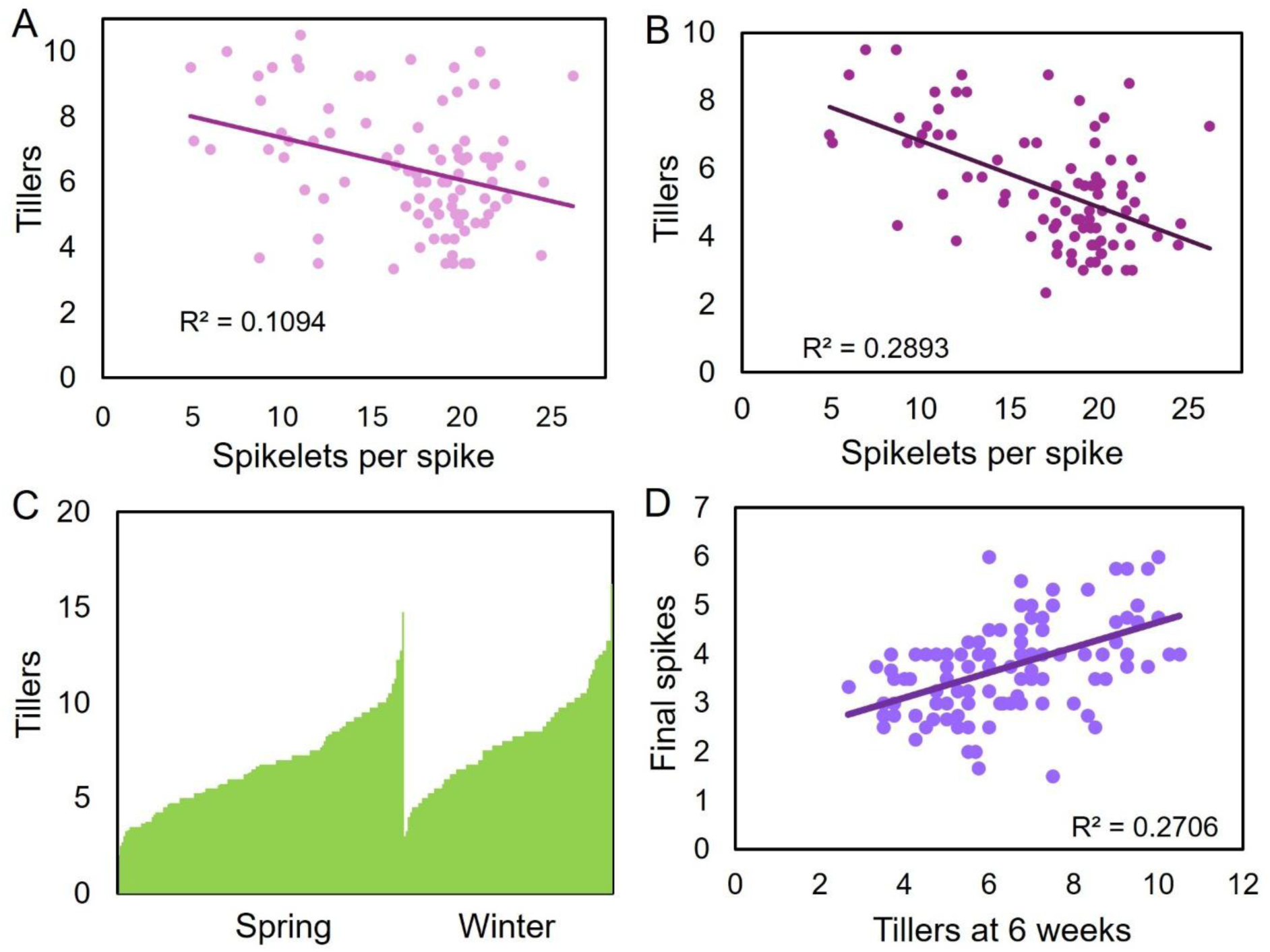
Negative correlation between spikelets per spike and tillers per in wheat landraces. **A-B)** Scatterplots showing correlation between mean number of tillers per plant and mean number of spikelets per spike after (A) 6 weeks of growth in 92 spring wheat landraces (n=3-5 per line, R^2^ = 0.11) and (B) 9 weeks of growth in 90 spring wheat landraces (n=3-5 per line, R^2^ = 0.29). Each point represents the mean values for an individual landrace **C)** Bar plot showing diversity in tiller number after 6 weeks of growth in a panel of 248 spring and winter landraces. **D)** Scatterplot showing correlation between mean number of spikes per plant and mean number of tillers after 6 weeks of growth in 114 spring wheat landraces (n=3-5 per line, R^2^ = 0.27). Each point represents the mean values for an individual landrace.

**Figure 2:**
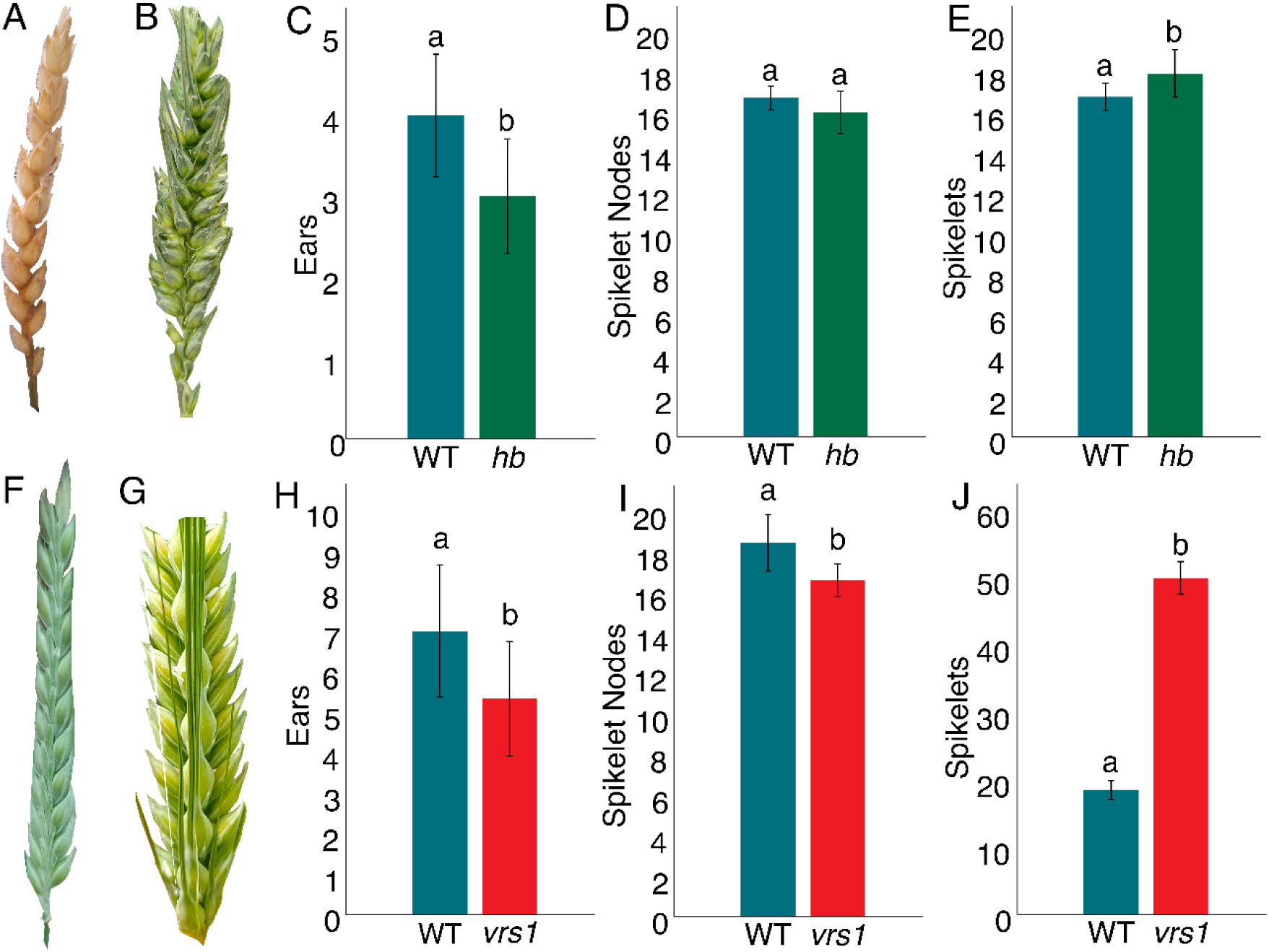
Tiller and spikelet development in *hb* wheat and *vrs1* barley. **A-C)** Measurements of reproductive architecture in the *hb* mutant compared to the WT parent line; mean final spike number (A), mean spikelet nodes per spike (B), mean spikelets per spike (C). Error bars are 1 standard deviation from the mean. Different letters above bars indicate statistically significant difference between lines (two-tailed t-test, P<0.05, n=7-8). **D-F)** Measurements of reproductive architecture in the *vrs1* mutant compared to the WT parent line; mean final spike number (A), mean spikelet nodes per spike (B), mean spikelets per spike (C). Error bars are 1 standard deviation from the mean. Different letters above bars indicate statistically significant difference between lines (two-tailed t-test, P<0.05, n=7-9).

This correlation is already present 6 weeks after germination (Supplementary Figure 1A), and strengthens by 9 and 12 weeks post germination (Figure 1, Supplementary Figure 1B), as a higher proportion of lines have undergone the floral transition. During the vegetative phase, the difference in tillering between wheat varieties is largely due to their inherently different rates of tillers (Supplementary Figure 1C). Tiller number at 6 weeks post germination is therefore a relatively poor predictor of final spike number (Supplementary Figure 1D). However, once flowering is initiated, the major tillers will form fully developed spikes and undergo stem elongation, whereas the other tillers will senescence to remobilise nutrients to the developing spikes. Tiller number at 12 weeks is therefore primarily a function of the number of spikes developing, rather than the initial rate of tillering. Our data suggest that plants with larger spikes (more spikelets per spike) sustain fewer tillers, and therefore ultimately make fewer spikes. This trade-off is indicative of a correlative inhibition effect, but these data do not suggest in which direction the inhibition acts (tillers inhibit spike size, or spike size inhibits tillering).

### High spikelet number wheat and barley lines produce fewer spikes

The data in Figure 1 only indicate a correlation between tillering and spikelets per spike. To determine if there is a causal link between these traits, we used mutants in wheat and barley that have increased spikelet number per spike, but which are otherwise near-isogenic with their parent lines. In wheat, the *highly branched (hb)* line has previously been reported to produce a high number of paired spikelets (Dixon *et al*., 2018), resulting in a significantly increased number of spikelets per spike over the wild type (Figure 1C). Likewise, the *vulgare-row spike/six-row spike1* (*vrs1*) mutant of barley produces a significant increase of spikelets per spike compared to its parent Bowman line (Figure 1F)(Druka et al, 2011). Both lines also produced significantly fewer tillers and spikes than their wild-type parent line (Figure 1A, D). These lines show the same apparent trade-off between spikelets per spike and reproductive tiller number/spike number observed among landraces, and in turn suggest that the correlative inhibition occurs from the developing spikelets in existing spikes towards the initiation of new tillers and spikes. These lines were therefore used to test the hypothesis that spikes of wheat and barley suppressed tillering in proportion to their spikelet number.

### Development of later-emerging spikes is inhibited in high spikelet lines

It is well established in cereals that the spike produced by the primary shoot meristem is the largest on the plant; each subsequent spike produced is smaller than the primary spike, roughly in proportion to its subsequent time of emergence (Vergara *et al*., 1990; Mohapatra and Kariali, 2008). This ‘productivity gap’ could simply be a function of the time each spike spends developing; however, it might also represent the effects of correlative inhibition. That is to say that, as well as inhibiting the formation of new tillers/spikes *per se*, the spikelets present in the early spikes might also inhibit the development (and therefore spikelet number) of later-initiating spikes. To test the hypothesis that the productivity gap observed in cereals is caused by correlative inhibition, we carefully analyzed the development of *hb* and *vrs1* spikes relative to their wild-type parents.

For barley, tillering was tracked from germination onwards, so that we knew the order of emergence of the tillers (‘ordinal tiller number’). At 35 days after germination, each tiller on the plant was dissected, and the number of spikelet ridges present on the developing spike was counted, and the relative stage of development was noted. When considered as whole plants, the mean number of spikelet ridges per spike does not differ between vrs1 and WT (Figure 3A), and nor does the mean stage of development of those spikes (Figure 3B). However, if we consider the data for each ordinal tiller number separately, a different pattern emerges. In terms of spikelet ridges, the first 10 tillers to emerge (include the main shoot) show no differences between *vrs1* and WT, but for the subsequent 3 tillers, the *vrs1* spikes have fewer spikelet ridges than the equivalent wild-type spike, significantly so in the case of the 13^th^ spike (Figure 3E). At this stage, wild-type plants have also initiated on average 6 more developing spikes than *vrs1* mutants (Figure 3E). A similar pattern is seen for stage of development, with the addition that the earliest spikes in *vrs1* are also further ahead in development than wild-type equivalent (statistically significantly for the main shoot). These data are consistent with the early spikes in *vrs1* exerting a stronger ‘dominance’ over both later-initiating spikes and over spike initiation *per se* than wild-type spikes, as a result of their increased spikelet number per spike.

**Figure 3:**
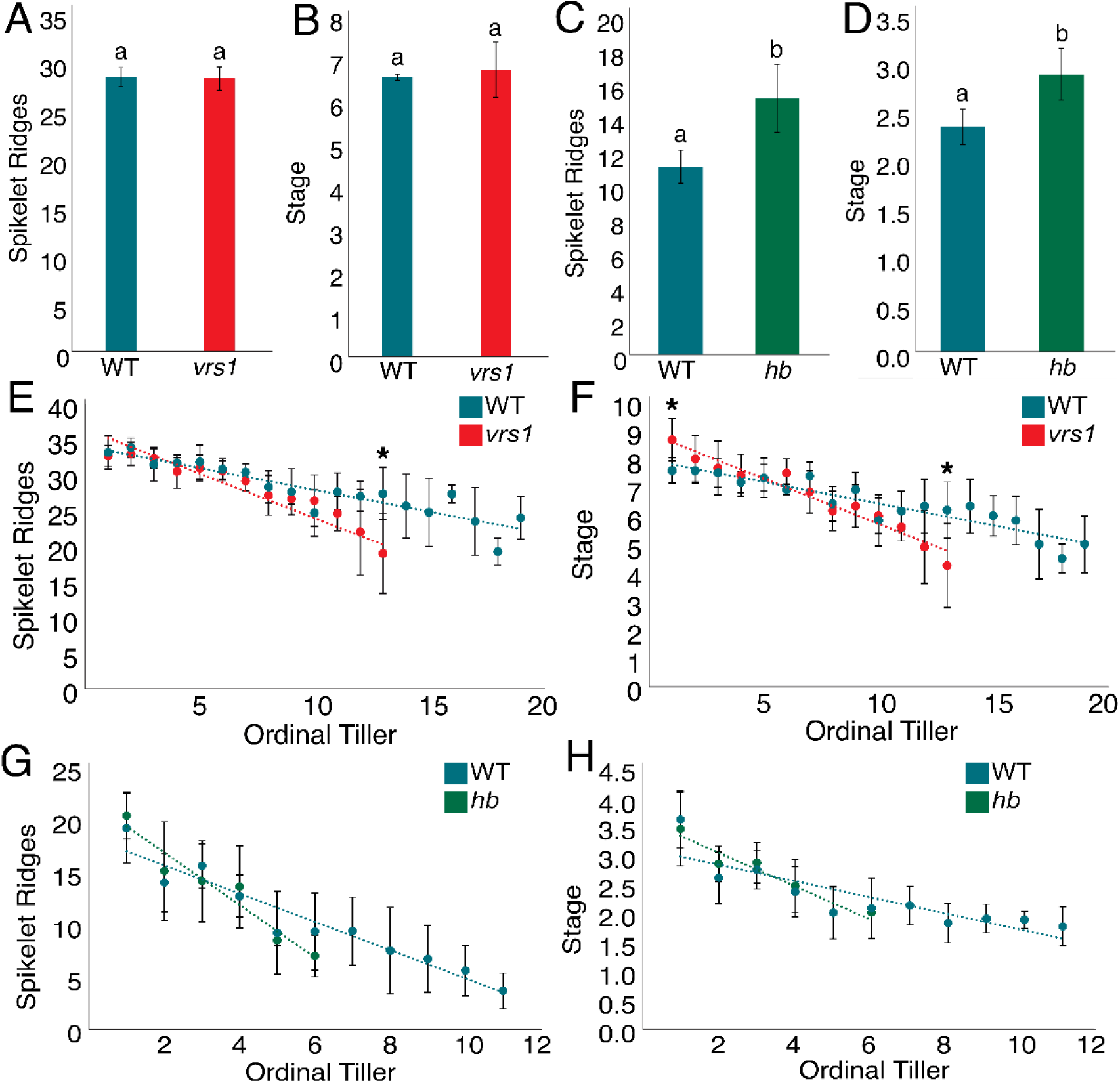
Analysis of SAM development in high spikelet lines. **A,B)** Graphs showing mean spikelet ridge number per spike per plant (A) and mean stage of spike development per plant (B) in *vrs1* and WT barley at 35dpg. Error bars are 1 standard deviation from the mean. Different letters above bars indicate statistically significant difference between lines (two-tailed t-test, P<0.05, n=6-7). **C,D)** Graphs showing mean spikelet ridge number per spike per plant (A) and mean stage of spike development per plant (B) in *hb* and WT wheat at 42dpg. Error bars are 1 standard deviation from the mean. Different letters above bars indicate statistically significant difference between lines (two-tailed t-test, P<0.05, n=10). **E,F)** Graphs showing mean spikelet ridge number (A) and mean stage of development for each ordinal tiller separately, in *vrs1* and WT barley at 35dpg. Error bars are 1 standard deviation from the mean. An asterisk above a point denotes statistically significant difference between lines at that ordinal position (two-tailed t-test, P<0.05, n=6-7). **G,H)** Graphs showing mean spikelet ridge number (A) and mean stage of development for each ordinal tiller separately, in *hb* and WT wheat at 42dpg. Error bars are 1 standard deviation from the mean. An asterisk above a point denotes statistically significant difference between lines at that ordinal position (two-tailed t-test, P<0.05, n=10).

For wheat, we performed the same analyses, with dissections at 42 days post germination. At this stage, *hb* had clearly initiated fewer spikes than WT (Figure 3G), but there was no clear difference in the number of spikelet ridges per spike in later spikes, nor in the stage of development of later spikes (Figure 3G,H). Thus, in *hb* wheat the effect of more developing spikelets per spike in the earlier spikes is largely manifested as reduction in overall spike number, rather than a productivity gap between spikes that do initiate.

Collectively, these results do not strongly suggest that the inherent productivity gap between successively-initiated tillers is primarily driven by correlative inhibition, although they do suggest that in later-initiating spikes, the effect is exaggerated by correlative inhibition from the earlier spikes.

### Ablation of spikelets causes proportional increases in tillering

To formally test for the existence of spikelet-driven feedback on tillering/spike initation, we conducted a series of experiments in which certain shoot meristems were surgically ablated. We hypothesized that, by removing the hypothesized repressive effect of spikelets, the ablation of meristems would result in increased tillering and spike initiation in the treated plants. Furthermore, we hypothesized that this tillering increase would be proportional to the number of spikelets ablated.

Firstly, we ablated the main shoot meristem in *vrs1* and wild-type barley, at a time point (28 dpg) where the reproductive transition has occurred, and spikelets are being initiated. This ablation resulted in a significant increase in both the rate of tillering, and in the maximum tiller number in both *vrs1* and wild-type (Figure 4A). Indeed, tillering in the ablated *vrs1* plants now matched the tillering observed in untreated wild-type. Furthermore, the effect was proportionally greater in the *vrs1* plants (a 13% and 26% increase in maximum tiller number resulting from ablation in bowman and *vrs1* respectively). These data strongly support the idea that early, ‘dominant’ spikes can suppress tiller initiation.

**Figure 4:**
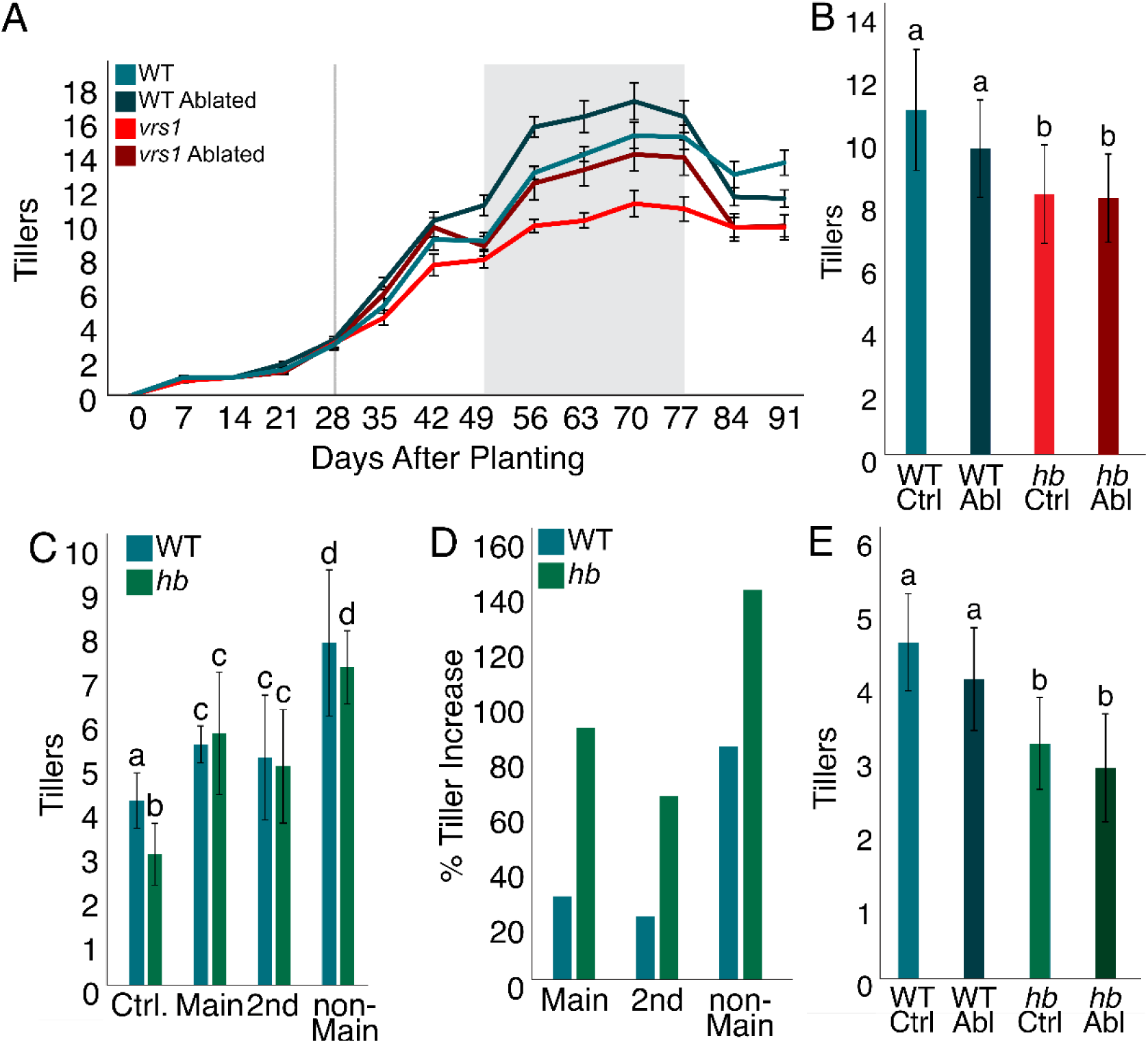
The effect of SAM ablation on barley and wheat. **A)** Mean tiller number per plant over time in WT and vrs1 plants with either the main meristem ablated or left intact. The point of ablation is represented by the grey vertical line at 28 Dpg. From 49 to 77 dpg (shown in grey) *vrs1* mean tiller number was significantly less than *vrs1* ablated and WT, both of which were not significantly different from each other, but produced significantly fewer tillers than ablated WT (ANOVA, P<0.05, n=10). **B)** Graph showing mean maximum tiller number per plant (63dpg) in WT and *vrs1* barley plants either left intact or with the main shoot meristem ablated at 42 dpg. Error bars are 1 standard deviation from the mean. Bars with the same letter are not statistically different from each other (ANOVA, P<0.05, n=8-10). **C,D)** Graphs showing mean maximum tiller number per plant (63 dpg) in WT and *hb* wheat plants either left intact (Ctrl), or with ablation of the main shoot meristem (main), the meristem of the first initiated tiller (1^st^), or all meristems besides the main shoot (non-main) at 28 Dpg (C), and percentage increase in maximum tiller number relative to control plants (D). Error bars are 1 standard deviation from the mean. Bars with the same letter are not statistically different from each other (ANOVA, P<0.05, n=9-10). **E)** Graph showing mean maximum tiller number per plant (56dpg) in WT and *hb* wheat plants either left intact or with the main shoot meristem ablated at 42 dpg. Error bars are 1 standard deviation from the mean. Bars with the same letter are not statistically different from each other (ANOVA, P<0.05, n=8-10).

A more complex set of expanded ablations were carried out in wheat, in which either 1) the main shoot meristem, 2) the meristem of the first initiated tiller or 3) every tiller except the main shoot, were ablated in both wild-type and *hb* at 28 dpg. As in barley, ablation of meristems resulted in an increase in tillering in both wild-type and *hb*, for all treatments (Figure 4C). The observed effect was stronger in *hb* than wild-type, and was also proportional to the number of spikelets ablated (Figure 4D). Thus, ablation of the main meristem resulted in a stronger response than ablation of the 1^st^ tiller meristem, while ablation of all non-main meristems resulted in an even stronger response.

These ablations were conducted on meristems actively generating spikelets. To test whether the effect is driven by the presence of spikelets *per se*, or by the active development of spikelets, we conducted a further set of ablations on both barley and wheat plants at 42 dpg, when main shoot SAMs had reached the terminal spikelet stage, at which point shoot meristems are no longer actively generating new spikelets. Unlike the ablation of actively developing spikelets, these ablations resulted in no significant increase in tiller number in any species or line (Figures 4B and 4E). Taken together these data strongly support the hypothesis that a feedback mechanism influences tiller development that is dependent specifically on the number of actively developing spikelets.

### Cytokinin treatment increases tiller number and spikelet number

Correlative inhibition phenomena in plants have long been associated with phytohormone signalling. We therefore hypothesized that the observed correlative inhibition between spikelets and tiller initiation in cereals is driven by such a mechanism. More specifically, we hypothesized that the effect is driven by developing spikelets/spikes acting as a sink for *trans*-Zeatin cytokinin from the root system and thereby limiting the pool of *trans*-Zeatin available to other meristems (Walker *et al*., 2023). Cytokinin has long been associated with branch initiation/outgrowth (Walker and Bennett, 2024) and with the rate of meristematic activity in inflorescences (Landrein *et al*., 2018; Walker and Bennett, 2024), and it would therefore be plausible that decreasing availability of *trans*-zeatin would prevent new tiller initiation and affect meristem size in later-initiating spikes.

To test this hypothesis, we first developed a treatment system based on a previously described injection system (Li *et al*., 2021) in which individual wheat tillers of the elite spring wheat Cadenza were injected with the synthetic cytokinin 6-Benzylaminopurine (6-BA) (see methods). We performed this treatment for either 0 weeks (control), 2 weeks (END1), 4 weeks (END2) or 6 weeks (END3) beginning at 21dpg, and measured the effects on meristem and tiller development. Consistent with many previous observations, 6-BA treatment increased tiller number compared to untreated controls between 35 and 56dpg, with a stronger increase with longer treatment durations (Figure 5A). No significant change in number of spikelet nodes per spike at 49dpg was observed (Figure 5C), however a significant increase in paired spikelet formation was observed, commensurate with the length of time the shoot system was exposed to increased cytokinin (Figure 5D). Unexpectedly, we also saw a decrease in the rate of meristem development at 49dpg in response to the cytokinin treatment (Figure 5B). All treated plants were in an earlier stage of development than untreated controls, and plants with extended (4 week) cytokinin treatments were significantly delayed even compared to shorter (2 week) treatment plants. Taken together, these data demonstrate that cytokinin supply affects both tiller initiation and meristem development in wheat.

**Figure 5:**
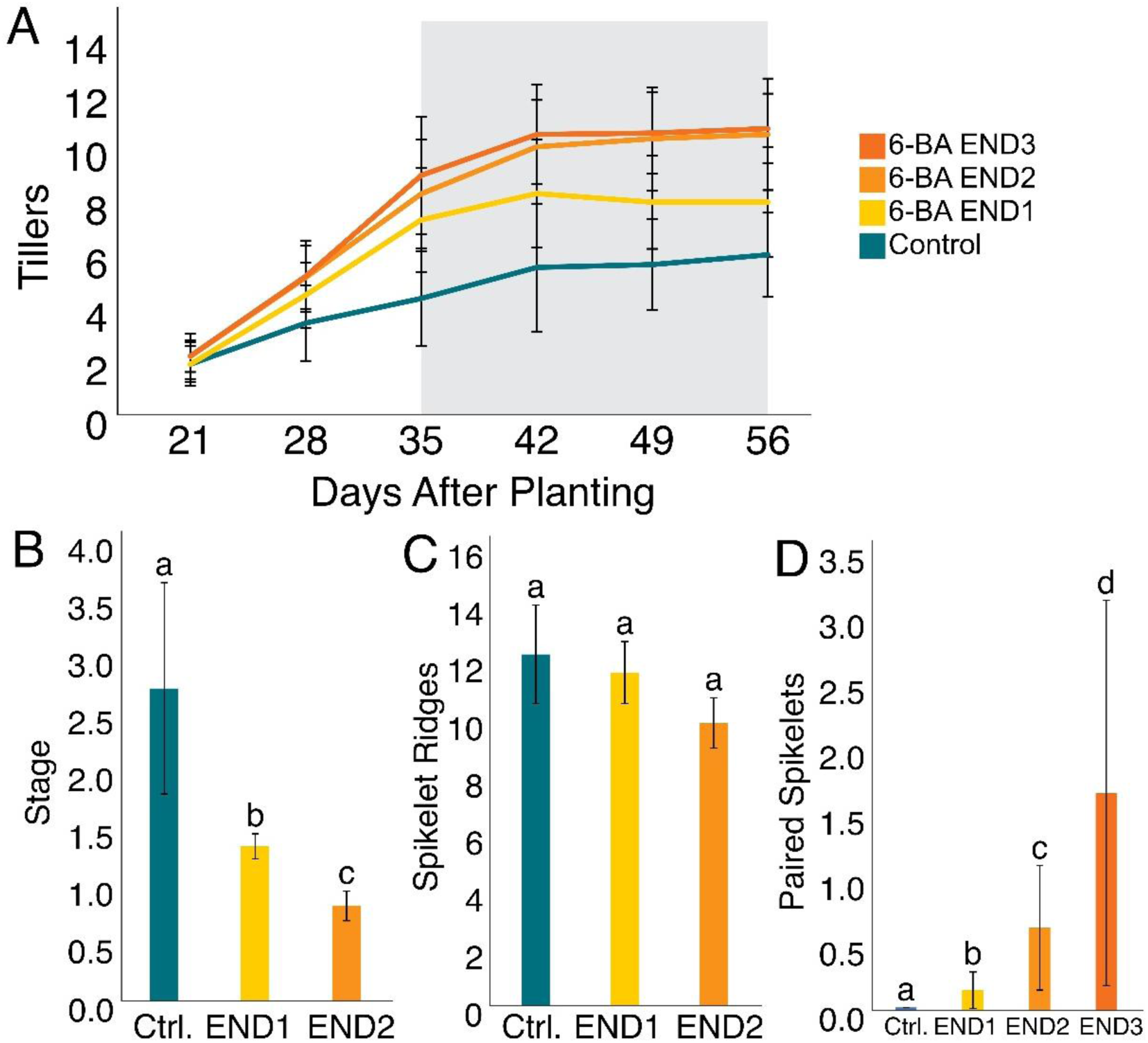
The effect of cytokinin on wheat shoot development. Plants were treated weekly with 0.33ml of 100μM synthetic cytokinin 6-BA from 21 Dpg, for 2 weeks (END1); 4 weeks (END2); 6 weeks (END3), or no weeks (Control/Ctrl). **A)** Graph showing mean number of tillers per plant over time. Between 35 and 56 Dpg, the grey background indicates statistically significant difference between Control, END1 and the END2/END3 treatments (which are not significantly different to each other)(ANOVA, P<0.05, n=12-16). **B,C)** Graphs showing the mean stage of meristem development (B) and mean number of spikelet ridges in control, END1 and END2 plants at 49dpg. Error bars are 1 standard deviation from the mean. Bars with the same letter are not statistically different from each other (two-tailed t-test, P<0.05, n=3-4). Dissections were made after 4 weeks of treatment, therefore at this point, there is no distinction between the END2 and END3 treatments. **D)** Graph showing the mean number of paired spikelets produced per spike in control, END1, END2 and END3 treatments. Data taken at end of plant life. Error bars are 1 standard deviation from the mean. Bars with the same letter are not statistically different from each other (ANOVA, P<0.05, n=11-14).

### Cytokinin delays reproductive development in wheat

To further investigate the apparent effect of cytokinin on the rate of meristem development, a rapid-flowering wheat landrace (CIMMYT CWI 7129) was treated with or without 6-BA throughout its development beginning at 21dpg. During this experiment, 3-4 plants were dissected every 48 hours, to determine the mean stage of meristem development. Plants transitioned to reproductive development after 24dpg, and after this point the cytokinin-treated plants were significantly retarded in meristem development compared to the control plants (Figure 6). This gap increased over the course of the experiment and by 38dpg the control plants (Waddington stage 8.0) were in a much more advanced stage than the 6-BA treated plants (Waddington stage 4.0). Alongside the experiments reported in Figure 5, this data strongly supports the conclusion that cytokinin has a previously unreported influence on the rate of meristem development in wheat.

**Figure 6:**
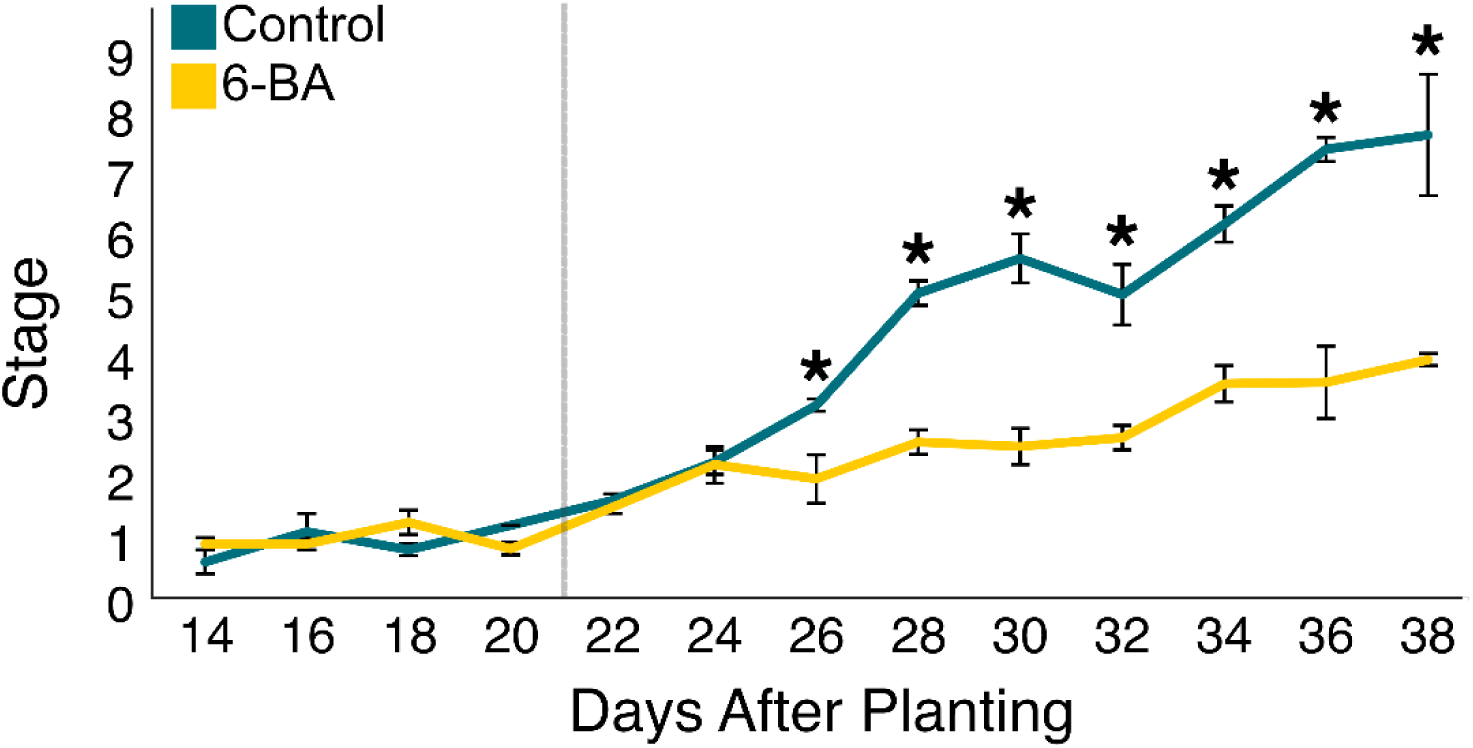
The effect of cytokinin on rate of SAM development. Graph showing mean developmental stage in wheat plants treated with or without a weekly application of 0.33ml of 100μM synthetic cytokinin 6-BA after 21 dpg (indicated by the grey line), until the end of the experiment. Error bars are 1 standard deviation from the mean. An asterisk indicates a statistically significant difference between treatments at that time point (two-tailed t-test, P<0.05, n=3-4).

### Cytokinin induced delay in spikelet production allows for an increase in tillering

Tiller and spikelet development was also tracked in this time-course experiment, to elucidate the role of cytokinin in spikelet-tiller feedback. The cytokinin treatment not only resulted in a reduction in the rate of meristem development, but in a reduction in the number of spikelet ridges initiating in each spike present on the plant (Figure 7B), which was visible after the first treatment, and at most timepoints up to 38dpg. Thus, the retardation of meristem development caused by cytokinin is associated – directly or indirectly – with a temporary reduction in the number of developing spikelets per spike, early in reproductive development. This further corresponded with a significant increase in the number of tillers produced by the plants that was observed from 28dpg onwards (Figure 7A). While we initially hypothesized that increased cytokinin supply would, as a direct consequence, allow increased tiller initation, these data raise an alternative, non-mutually exclusive possibility. Namely, that increased cytokinin supply inhibits meristem development and spikelet initiation, thereby removing correlative inhibition upon tiller initiation that occurs through some other mechanism.

**Figure 7:**
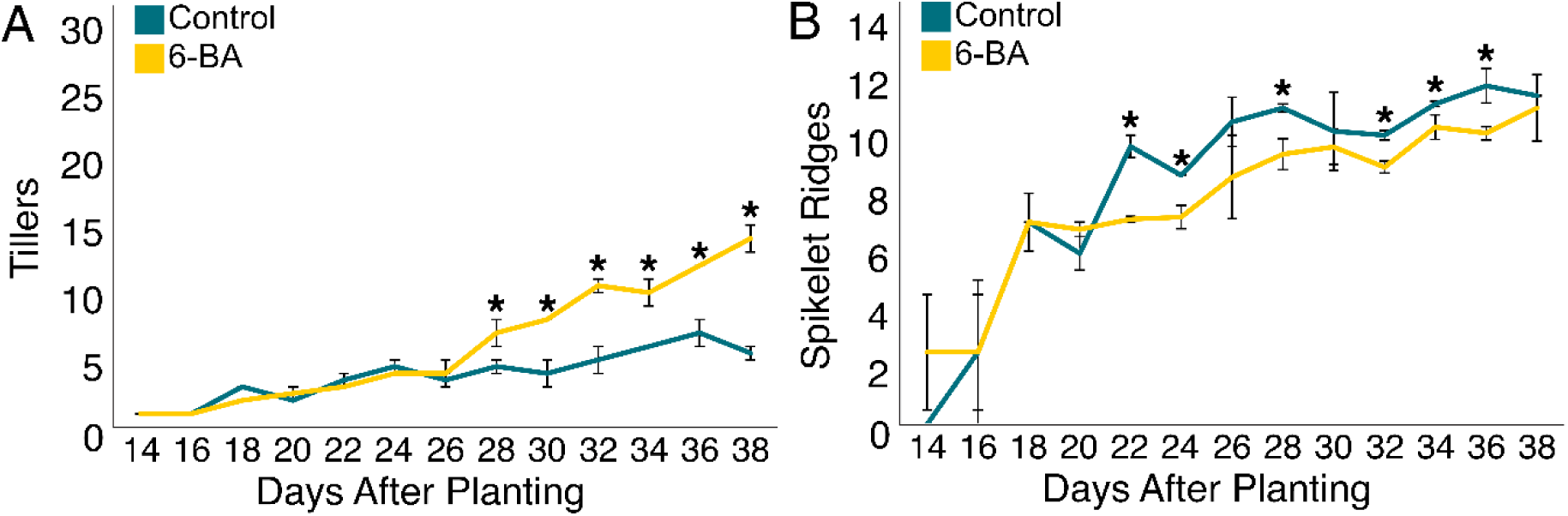
The effect of CK on tiller and spikelet development in wheat. A,B) Graphs showing the mean number of spikelet ridges per meristem (A) and mean number of tillers per plant (B) over time in wheat plants treated with or without a weekly application of 0.33ml of 100μM synthetic cytokinin 6-BA after 21 dpg until the end of the experiment. Error bars are 1 standard deviation from the mean. An asterisk indicates statistically significant difference between treatments at that time point (two-tailed t-test, P<0.05, n=3-4).

### Cytokinin overcomes spikelet-driven feedback in high spikelet lines

Finally, we attempted to test whether the enhanced spikelet-driven correlative inhibition observed in *vrs1* and *hb* mutants can be alleviated by cytokinin treatment. In wheat, both *hb* and wild-type plants were treated with or without 6-BA for 4 weeks from 21dpg. These treatments resulted in a significant increase in tillering in both lines, but the *hb* plants responded more strongly to the treatment than wild-type, resulting in treated *hb* plants producing the same number of tillers as treated wild-type plants (Figure 8A,B). We also observed a reduction in the mean developmental stage of meristems at 49dpg in both lines in response to cytokinin treatment, with the *hb* line again showing a stronger response (Figure 8C,D). The treatments did not show any effect on spikelet node number (Figure 8E) but did cause increases in paired spikelet formation (Figure 8F). Cytokinin treatment resulted in wild type plants producing the same number of paired spikelets as *hb* (Figure 8F).

**Figure 8:**
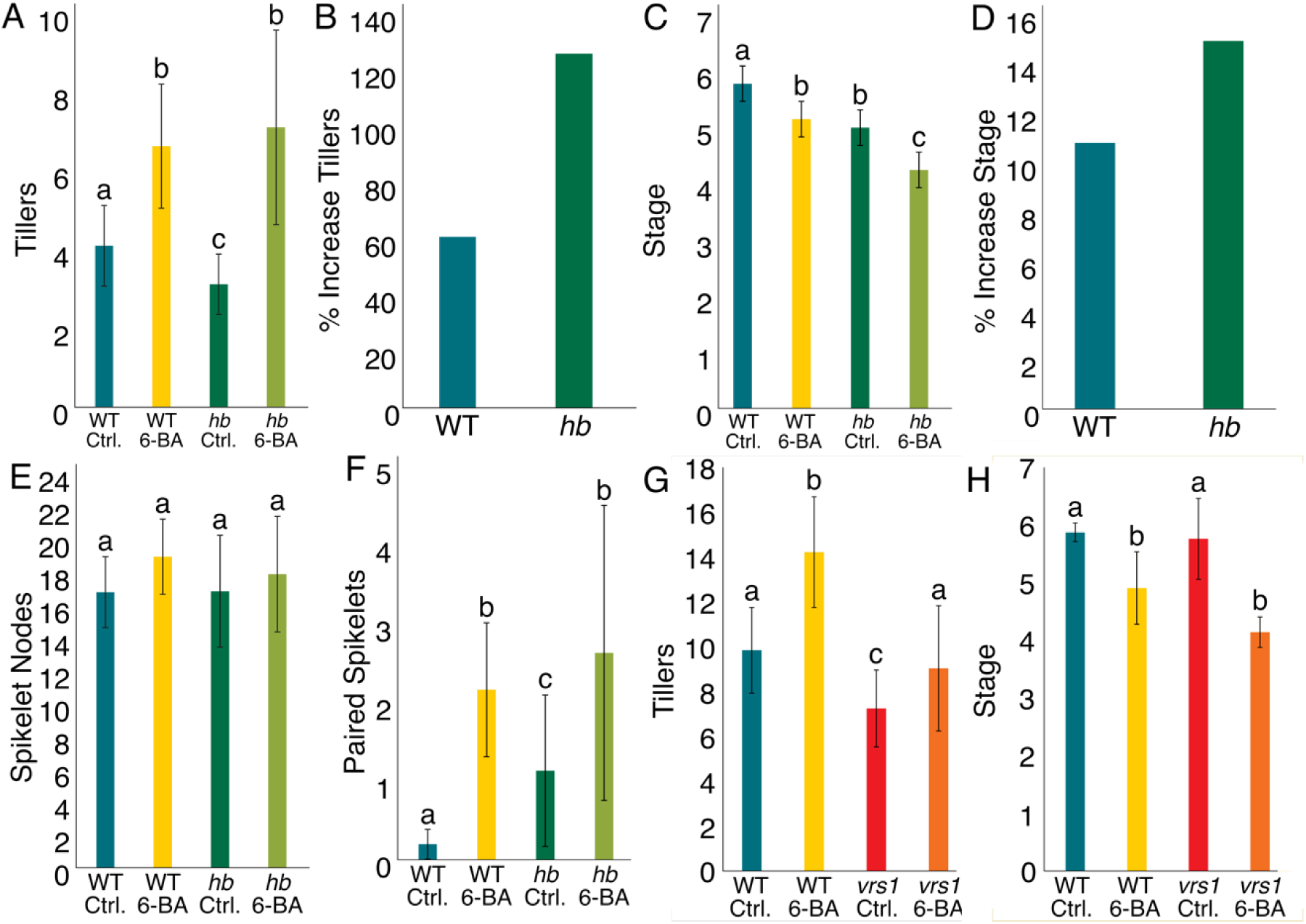
The effect of cytokinin on development in high spikelet lines of wheat and barley. **A,B)** Graphs showing mean maximum tiller number per plant (49 Dpg) in WT and hb wheat plants treated with or without a weekly application of 0.33ml of 100μM synthetic cytokinin 6-BA after 21 dpg for 4 weeks (A), and the percentage increase in maximum tiller number relative to untreated control plants in both lines (B). Error bars are 1 standard deviation from the mean. Bars with the same letter are not statistically different from each other (ANOVA, P<0.05, n=13-15). **C,D)** Graphs showing mean meristem developmental stage (49 Dpg) in WT and hb wheat plants treated with or without a weekly application of 0.33ml of 100μM synthetic cytokinin 6-BA after 21 dpg for 4 weeks (A), and the percentage decrease in developmental stage relative to untreated control plants in both lines (B). Error bars are 1 standard deviation from the mean. Bars with the same letter are not statistically different from each other (ANOVA, P<0.05, n=3-4). **E,F)** Graphs showing mean spikelet node number per spike at 49dpg (E) and number of paired spikelets at 49 Dpg (F) in WT and hb wheat plants treated with or without a weekly application of 0.33ml of 100μM synthetic cytokinin 6-BA after 21 dpg for 4 weeks. Error bars are 1 standard deviation from the mean. Bars with the same letter are not statistically different from each other (ANOVA, P<0.05, n=3-4). **G,H)** Graphs showing maximum tiller number per plant (56 Dpg)(A) and mean meristem developmental stage at 49 Dpg (B) in WT and *vrs1* barley plants treated with or without a weekly application of 0.33ml of 100μM synthetic cytokinin 6-BA after 21 dpg for 4 weeks. Error bars are 1 standard deviation from the mean. Bars with the same letter are not statistically different from each other (ANOVA, P<0.05, (G) n=12-13 (H) n=3-4).

In barley, *vrs1* and wild-type plants were also treated with or without 6-BA for 4 weeks from 21dpg. As was observed in wheat, the cytokinin treatment resulted in increased tillering in both lines (Figure 8G), and a delay in the mean developmental stage of meristems at 42dpg was also observed in both lines (Figure 8H). Our results in both wheat and barley are thus consistent with the hypothesis that cytokinin plays a key role in the observed correlative inhibition between developing spikelets and tiller/spike initiation.

## DISCUSSION

The results presented here indicate that a form of correlative inhibition plays a key role in shaping reproductive architecture in wheat and barley, and may explain some of the previously observed yield trade-offs in these species. We have shown that developing spikelets act as a source of ‘dominance’ and are able to inhibit the initiation of new tillers, and the development of further spike meristems, in both species (Figure 4). The effect exerted by the developing spikelets is topologically similar to apical dominance, in the sense that the development of new axillary meristems is inhibited by the cumulative ‘spikelet load’. However, since the dominance is exerted by secondary inflorescences, and those which are developing rather than fully active, there are also key differences to apical dominance. Although (Bangerth, 1989) inferred the existence of such correlative inhibition effects in grasses, and the grass phenology certainly supports the existence of such feedbacks (Dixon *et al*., 2018; Zwirek, Waugh and McKim, 2019; Sadka *et al*., 2023), there has been very little study of these phenomena in grasses. Our study thus unites cereal crop research with well-established research into correlative inhibition in both fruit crops and model species such as Arabidopsis (Sadka *et al*., 2023).

Our results also show that the availability of cytokinin plays a key role in moderating this correlative inhibition, promoting the development of tillers, and overcoming the dominant effect of increased spikelet load (Figure 8). The data from these experiments revealed previously unreported developmental roles for cytokinin in wheat. Firstly, cytokinin delays shoot meristem ageing (Figures 5B and 6). Secondly, cytokinin concentration results in proportional increases in the frequency of paired spikelet formation, without significantly affecting spikelet node number (Figure 5C-D). We propose that the delay in meristem development results in increased tillering by delaying the establishment of the full spikelet load-derived correlative inhibition on tillering. Cytokinin has long been associated with the promotion of branching, although the direct effects of cytokinin on branching are generally mild and somewhat enigmatic (Walker and Bennett, 2024). While cytokinin can certainly down-regulate expression of the branching regulatory gene *BRANCHED1* in Arabidopsis and pea axillary buds (Dun *et al*., 2012), the consequences of this down-regulation might be better viewed as priming buds for activation, rather than necessarily driving their outgrowth (Seale, Bennett and Leyser, 2017; Walker and Bennett, 2018). Our results are thus consistent with a more nuanced role for cytokinin in the control of branching, one acting by modulating the inhibitory strength of different shoot organs, rather than directly causing bud outgrowth. An alternative, non-mutually exclusive possibility is that shoot organs compete for the availability of root-derived cytokinin to drive their growth (Walker *et al*., 2023). If there is a relatively constant pool of root-derived *trans*-Zeatin (*t*Z) cytokinin during reproductive development, then as the number of cytokinin sinks in shoot system increases over time, the availability of *t*Z might eventually become too low to sustain the growth of any new shoot axes. However, although this ‘cytokinin dilution’ model is appealingly simple, it does not appear consistent with the effects observed here, in which spikelets are dominant when developing, but not when mature. The dominance of spikelets seems a more active phenomenon, rather than a passive one based on sequestering cytokinin.

It is also likely that other phytohormonal signals are involved in the observed effects. In particular, auxin export from dominant organs has long been associated with their ability to inhibit the development of other organs (Bangerth, 1989; Walker and Bennett, 2018; Sadka *et al*., 2023). One model to explain the effects of auxin is based on the canalization of auxin transport between organs (Prusinkiewicz *et al*., 2009), ultimately deriving from the canalization model of vascular patterning in plants (Sachs, 1981). Canalization models suppose that the growth of a new organ is connected to its ability to export auxin, which requires the formation of a canalized auxin transport link to an auxin sink in the connecting stem; the ability of this canalized link to form is dependent on both the auxin source strength of the new organ, and the auxin sink strength of the stem. Thus, it is hypothesized that the auxin exported from dominant organs inhibits new organ growth by weakening the auxin sink strength of the shared stem and preventing new canalized links from forming (Prusinkiewicz *et al*., 2009). In this context, the role of cytokinin in delaying spikelet development might be conceived as delaying the establishment of peak auxin export from early-formed spikes, thereby giving a greater window in which later-forming spikes can form canalized links to the shared stem, and begin their outgrowth. However, while such models are theoretically supported and can explain apparently self-organizing patterns of shoot development, they are also difficult to directly test experimentally and remain open to criticism (Krupinski and Jonsson, 2010; Ravichandran, Linh and Scarpella, 2020).

The results presented here are highly relevant to the aim of creating future cereals that produce increased grain yields without needing to increase resource availability (Alexandratos and Bruinsma, 2012; DeLucia *et al*., 2014). We provide evidence that correlative inhibition mechanisms are actively restraining yield potential, resulting in a negative correlation of individual spike productivity with spike number, and hindering the development of lines that produce a large number of highly productive spikes. Our results demonstrate that the production of spikes is not inherently limited by resource availability, and can be increased by a simple hormonal treatment. Thus, by further investigation of the role of cytokinin and other hormonal signals on correlative inhibition mechanisms, it may be possible to boost cereal yields in a way that does not require increased resource use. Tuning of cytokinin levels in specific tissues and time points has already been proposed as a promising pathway for improving crop yields (Chen *et al*., 2020; Rathore *et al*., 2024). The ability of cytokinin to regulate the rate of meristem development represents one clear target for yield improvement, if it allows the formation of more shoot axes. Recent improvements in our understanding of the roles of phytohormones in cereal development (Awale and McSteen, 2023; Bai *et al*., 2024; Szala *et al*., 2024; Walker and Bennett, 2024), and the increased availability of tools available for this research (Isoda *et al*., 2021; Zhang *et al*., 2022), thus hold the promise of unlocking yield improvements in our most important crops.

## MATERIALS & METHODS

### Plant material

For figures 2, 3, 4, 5 and 8, we used lines of spring barley (*Hordeum vulgare*) and hexaploid spring wheat (*Triticum aestivum*). The elite spring wheat Cadenza and the elite spring barley Bowman were used as wild-type lines in this work. The *hb* (highly-branched) wheat mutant in the Cadenza background was a kind gift of Laura Dixon (University of Leeds). The *vrs1* barley mutant was provided by Sarah McKim (University of Dundee) and the Barley Genetics group at the James Hutton Institute.

For Figures 1, 6 and 7, we used the YoGI wheat diversity panel, a kind gift of Andrea Harper (University of York)(Barratt et al, 2023). Figures 6 & 7 utilised CIMMYT CWI 7129/YoGI 028.

### Plant growth

All plants were grown in John Innes No. 2 compost. The day/night cycle was 16h/8h of light and dark at 20°C and 16°C respectively. Plants were grown in greenhouses with a natural light source alongside LED lights with an average light intensity of ∼250 µmolm^-2^s^-1^. Plants were grown in 500ml soil volume.

### Architecture measurements

Tiller counts were taken by measuring the number of emerged, distinct shoots (tillers) on the plant. Tillers were counted as emerged, if more than 50% of the tiller is visible from the leaf that ensheathes it. Spike counts were taken as a measurement of the number of productive spikes that have fully emerged at the end of plant life. Unproductive spikes were defined by producing no seed, or remaining green when other spikes on the plant had completed seed filling. Spikelet counts were taken as a measurement of every fully formed spikelet along the rachis of the spike, at the end of plant life. Counts of paired spikelets included spikelets formed directly below the main spikelet at a rachis node.

### Cytokinin treatment system

45.05g of powdered 6-Benzylaminopurine (Sigma-Aldrich) was diluted in 20ml of ethanol then mixed with 180ml of distilled water to create 200ml of 1mM 6-BA. This stock was stored at -20°C. 100µM aliquots of 6-BA was created from this stock using serial dilutions up to one hour before treatment was conducted. 0.33ml of 100µM 6-BA was injected every 7 days into each fully emerged tiller. Injections were performed using a 0.5mm diameter needle attached to a 2ml syringe. The needle was angled at approximately 15 degrees from the tiller and inserted approximately 10mm above the predicted location of the shoot apical meristem. Control plants were injected with an equivalent dilution of ethanol in water using the same method at the same time. This protocol was first adapted from a previously described 6-BA treatment protocol (Li *et al*., 2021).

### Microscopy and meristem dissection

Meristem dissections were primarily performed on an Olympus SZ51 stereo light microscope at 2-40x magnification, lit using an Olympus KL 300 LED light source. Some dissections and imaging were performed using a Keyence VHX-7000 digital microscope. Cereals were removed from their growing medium and the shoot system separated from the root system. Each tiller was dissected using a FEATHER incision micro scalpel to reveal the shoot apical meristem. The stage of development was identified using the criteria outlined in (Kirby and Appleyard, 1987).

Transition between vegetative and reproductive development in the SAM was defined by the transition between the vegetative and double ridge stage for both species, as defined in (Kirby and Appleyard, 1987), wherein floral primordia form above the leaf primordia along the apex of the SAM, creating the distinct “double ridge” appearance and indicating that the apex now can produce reproductive tissue. Shoot apical meristems which had reached the double ridge stage or later were defined by these criteria as having initiated reproductive development.

Stage of development was determined and assigned a quantitative value according to the Waddington scale (Waddington, 1983), a quantitative measurement commonly used to measure spike development in wheat and barley using a scale of numbers from 0 (seedling emergence) to 10 (pollination). This scale allows each point in development to be assigned a quantitative value, allowing for statistical analysis and comparison not possible with phenotypic data.

Spikelet ridges were counted as all clearly defined spikelet ridges, based on the criteria outlined in Kirby and Appleyard (1987).

## ACKNOWLEDGEMENTS

We thank members of the Bennett lab for maintaining the wheat plants in Figure 1 during the hard year of 2019/2020, including Josie Driver and Mary McKay. We thank Sarah McKim for helpful discussions. We thank Sarah McKim, the University of Dundee, and the Barley Genetics group at the James Hutton Institute for providing germplasm, especially Pauline Smith, Richard Keith, Chris Warden, and Dr Joanne Russell.

## AUTHOR CONTRIBUTIONS

A.W. performed experiments and analyzed the data. A.W. and T.B. designed the study and wrote the manuscript.

## CONFLICT OF INTEREST

The authors declare they have no conflict of interest.

## FUNDING

AW is supported by the Natural Environment Research Council (NERC) Panorama DTP (NE/S007458/1). TB is supported by Biotechnology and Biological Sciences Research Council (BSSRC) grant BB/X001423/1.

## DATA AVAILABILITY

All figures in this manuscript are associated with raw data. All raw data will be made available upon request to the corresponding author.

## Notes

### Competing Interest Statement

The authors have declared no competing interest.

### Summary of Updates

Updated references and acknowledgements to correct attribute germplasm.

